# DeltaMut: An Integrative Database of AlphaFold2-Derived Missense Variant Structures

**DOI:** 10.64898/2026.06.03.729798

**Authors:** Erda Qorri, Krisztián Ádám, Bertalan Takács, Sára Shemesh, Dénes Buzafalvi, Valentin Varga, Emese Pekker, Lajos Pintér, Zoltán Hegedűs, Bernadett Csányi, Lajos Haracska

## Abstract

The widespread use of next-generation sequencing has led to a surge in the number of identified variants with uncertain effects on protein function. These variants pose a significant challenge in diagnostics and hinder patient treatment strategies. Numerous variant effect predictors (VEPs) are available to assess variant impact, but they primarily rely on sequence-derived information. The recent development of AlphaFold2 has raised questions about whether information retrieved from wild-type or predicted structures of missense variants can improve the predictive power of these algorithms. While the AlphaFold Protein Structure Database serves as a valuable resource for wild-type protein structures, a large-scale collection of missense variant structures is not available, limiting current efforts to wild-type conformations and a handful of modeled variants.

To address this limitation, we developed DeltaMut, a comprehensive database containing over 77,000 protein structures, including 65,000 pathogenic and neutral missense variants. All structural models were generated using ParaFold, a high-performance computing-optimized implementation of AlphaFold2.

The large-scale and systematic generation of variant protein structures distinguish DeltaMut as a unique resource for both expansive statistical studies and detailed, case-specific investigations of variant-induced structural changes. Furthermore, the DeltaMut database is freely accessible without registration.

**Highlights:**

- DeltaMut is currently the largest database of AlphaFold2-predicted variant structures.
- Contains 77,713 structures covering 12,101 wild-type and 65,612 variant proteins.
- 70.6% of predicted structures have high or very high confidence (pLDDT ≥ 70).
- Freely accessible web server with visualization and download of variant models.

## Introduction

Genetic variants play a pivotal role in the development of several pathological conditions and have been linked to increased susceptibility to several types of cancer (Reza et al. 2021). It is estimated that of the 4 million identified missense variants, only a small fraction has been clinically classified, leaving the majority as variants of uncertain significance (VUS) (Cheng et al. 2023). The number of reported VUS has surged in recent years, creating a paradox, where despite the wealth of novel information on genetic variants, limited understanding of their impact continues to hinder patient treatment strategies (Federici and Soddu 2020). As a result, the classification and reclassification of VUS remain a central challenge in precision medicine.

To address this emerging problem, numerous supervised and unsupervised machine learning prediction algorithms, referred to as VEPs, have been developed. Notable examples include AlphaMissense (Cheng et al. 2023), REVEL (Ioannidis et al. 2016), PolyPhen2 (Adzhubei, Jordan, and Sunyaev 2013), and EVE (Frazer et al. 2021). Most of these VEPs are predominantly trained on features derived from the amino acid sequence, such as evolutionary conservation (Ng and Henikoff 2002), while structural and physicochemical properties have historically been underutilized and restricted only to proteins with experimentally determined structures deposited in the Protein Data Bank (PDB) (Adzhubei, Jordan, and Sunyaev 2013), (Bromberg and Rost 2007).

Recent developments in protein structure prediction, particularly the introduction of AlphaFold2 (Jumper et al. 2021) and RoseTTAFold (Baek et al. 2021), have marked a transformative moment in VEP development by providing access to a wealth of structural information.

Moreover, the launch of the AlphaFold Protein Structure Database has made structural data easily accessible, encouraging the development of novel VEPs that incorporate features derived from AlphaFold2-predicted structures, such as AlphScore (Schmidt et al. 2023), MOVA (Hatano, Ishihara, and Onodera 2023), and QAFI (Ozkan, Padilla, and de la Cruz 2024). However, these efforts have largely concentrated on the wild-type native conformations of proteins. Structural features derived from the modeled structures of pathogenic and benign missense variants remain understudied despite recent reports suggesting that AlphaFold2 can capture local structural changes caused by mutations (McBride et al, 2023). Nonetheless, research leveraging features from mutated protein structures at scale remains limited.

A major limitation is the substantial computational and storage burden associated with predicting thousands of missense variant protein structures. To date, no comprehensive large-scale structural datasets of missense variants are publicly available, thus restricting the integration of structural information into variant classification frameworks. Furthermore, the absence of such comprehensive structural datasets hinders broader investigations into AlphaFold2’s behavior in predicting the structural impact of mutations.

To address this challenge, we present DeltaMut, a comprehensive database of predicted protein structures for thousands of pathogenic and neutral missense variants. In its first release, DeltaMut contains 77,713 protein structures modeled using ParaFold (Zhong et al. 2022), of which 12,101 are wild-type conformations ranging from 24 to 3,500 amino acids in length, while 65,612 are their corresponding missense variants sourced from Humsavar (Consortium 2024).

The DeltaMut web server is freely accessible without registration and provides a broad range of functionalities for its users. To our knowledge, DeltaMut represents the largest publicly available collection of modeled missense variant protein structures to date. We anticipate that this resource will support the development of novel structure-derived features for improving missense variant classification algorithms. In addition, DeltaMut’s extensive variant coverage can aid the design of targeted experimental studies by enabling researchers to visually inspect predicted structural disruptions of individual residues prior to experimental validation. The mutant protein structures, which can be freely downloaded from the webserver, may also serve as input data for studies modeling interactions between native and mutated protein structures.

## Materials and Methods

### Missense variant collection and processing

Missense variants used as input for the structural predictions were retrieved from Humsavar (https://www.uniprot.org/help?query=humsavar), a comprehensive repository of missense variants annotated in human UniProtKB/Swiss-Prot entries (release 2023_1; February 22, 2023) (“UniProt: The Universal Protein Knowledgebase in 2025” 2025). An initial dataset of 81,922 missense variants was downloaded and subjected to several rigorous filtering steps, such as excluding VUS, duplicates, and incomplete entries. The retained Humsavar variants were then screened against the ClinVar database (Landrum et al. 2014), and overlapping variants were discarded. Following these filtering steps, the final dataset utilized for structural predictions comprised 70,345 unique missense variants, (30,926 pathogenic and 39,419 benign), from 12,580 unique proteins (Figure 1).

**Figure 1:**
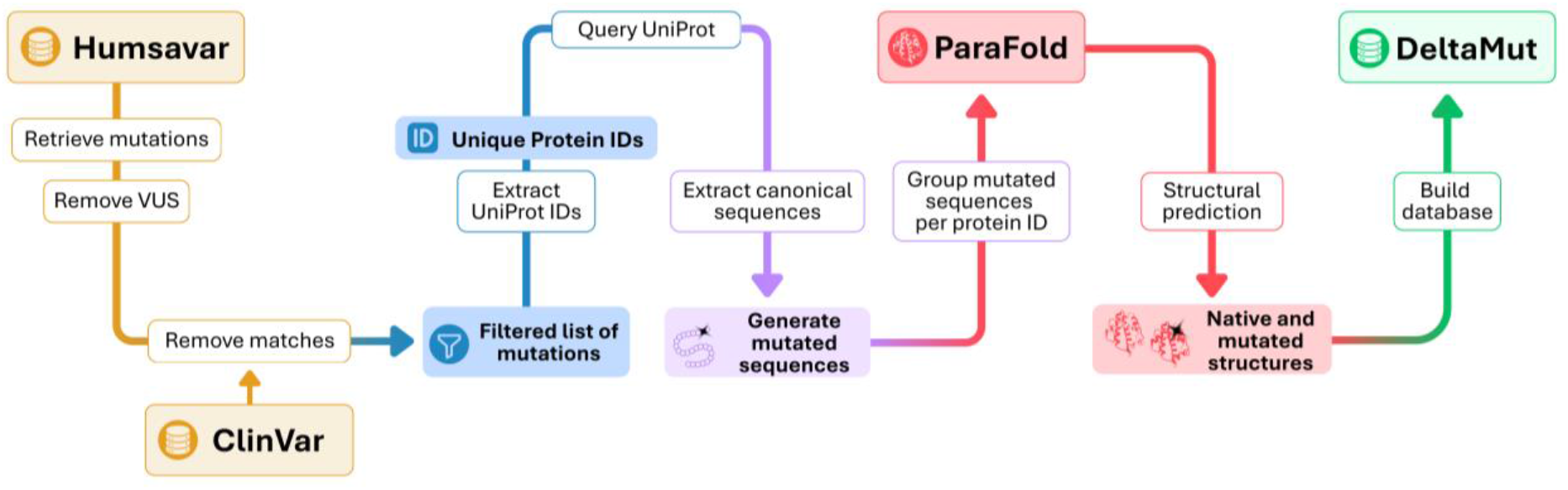
Workflow outlining structure prediction and DeltaMut development steps.

### Protein structure prediction

Structures of wild-type and mutated proteins were predicted using ParaFold (Zhong et al. 2022), a high-performance computing (HPC) implementation of AlphaFold2, optimized for large-scale protein structure predictions. All computations were performed on the *cpu, gpu*, and *bigdata* partitions of Komondor, Hungary’s largest HPC cluster (https://ncc.dkf.hu/en.html).

Similar to AlphaFold, ParaFold requires an amino acid sequence as input. In the case of wild-type structures, the canonical amino acid sequence was retrieved from the UniProt database (Consortium 2024). For the prediction of variant structures, the reference residue in the canonical amino acid sequence was substituted with its corresponding alternative residue.

The structure prediction step on the Komondor HPC was carried out in a two-stage process. Initially, predictions were submitted to the *cpu* partition to generate the *features*.*pkl* file, which contains the output of the multiple sequence alignment construction phase. The extracted features then served as input for model generation and structure refinement steps, carried out on Komondor’s *gpu* partition. Predictions that failed on either the *cpu* or *gpu* partitions were resubmitted to the more powerful *bigdata* partition of the Komondor HPC. ParaFold was run with default parameters as follows: *--model_present:* monomer_ptm; *--use_gpu:* True; *-database_present:* full_dbs; *--model_selection:* five models, *--recycling:* 3, and *–use_GPU_relax:* True.

### Structure prediction quality

The quality of the predicted structures reported in this study was assessed using the best-ranked relaxed model as reported in the *ranking_debug*.*json* file, which contains the ranked predicted models based on their mean predicted Local Distance Difference Test (pLDDT). The pLDDT score is a local confidence measure assigned to each predicted residue, ranging from 0 to 100, with higher scores indicating greater confidence in the predictions (Jumper et al. 2021). The mean pLDDT score of the best-ranked relaxed model was extracted for all structures: *wild-type, pathogenic*, and *benign* and utilized to classify these predictions into four confidence categories: very high confidence (pLDDT ≥ 90); high confidence (70 ≤ pLDDT < 90); low confidence (50 ≤ pLDDT < 70), and very low confidence (pLDDT <50).

### Database construction

The DeltaMut web server is built using the Elixir (Elixir Core Team, n.d.) and Phenix (Chris McCord, n.d.) frameworks. Variant data is loaded into an SQLite (Hipp 2020) database and augmented using metadata from the UniProt database (“UniProt: The Universal Protein Knowledgebase in 2025” 2025), including protein/gene names and UniProt IDs. PDB files are visualized using the Molstar package (Sehnal et al. 2021), with three preset modes: *Mutant model, Compare with wild-type model*, and *Mutant model confidence*. Protein visualization can be further configured through the built-in Molstar user interface. Structural superimposition of the mutated and wild-type structures is performed on demand by the web server without requiring further processing, using the *Superimposer* module from BioPython (Cock et al., 2009)

## Results and Discussion

### Prediction quality control

This release of DeltaMut includes a total of 77,713 protein structures of which 12,101 are wild-type and 65,612 are from pathogenic and benign missense variants. Among these, 29,523 structures correspond to pathogenic variants, while 36,089 represent benign ones. The length of the predicted proteins ranges from 24 to 3,487 amino acids, with 26.5% (17,396) of the 65,612 missense variant structures corresponding to proteins longer than 1000 amino acids. Notably, the dataset includes structures of missense variants identified in clinically relevant proteins associated with cancer (RASK, BRCA1, BRCA2, MLH1, MSH2, BARD1), neurodegenerative diseases such as Alzheimer’s (PSEN1, PSEN2), Huntington’s disease (HTT), and Parkinson’s disease (LRRK2, SNCA), as well as cardiovascular disorders including Long QT syndrome (CNQ1, KCNH2, SCN5A), Hypertrophic cardiomyopathy (MYH7, MYBPC3) and immunodeficiencies like Severe Combined Immune Deficiency (IL2RG).

Based on the pLDDT value of the best-ranked model, 70.6% (54,865) of the 77,713 structures were predicted with high or very high confidence, with 17.2% having pLDDT values above 90 and 53.4% scoring between 70 and 90. In contrast, 29.3% had average pLDDT values below 70, including 25% with pLDDT values between 50 and 70, and 4.23% with pLDDT <50 (Figure 2A).

**Figure 2:**
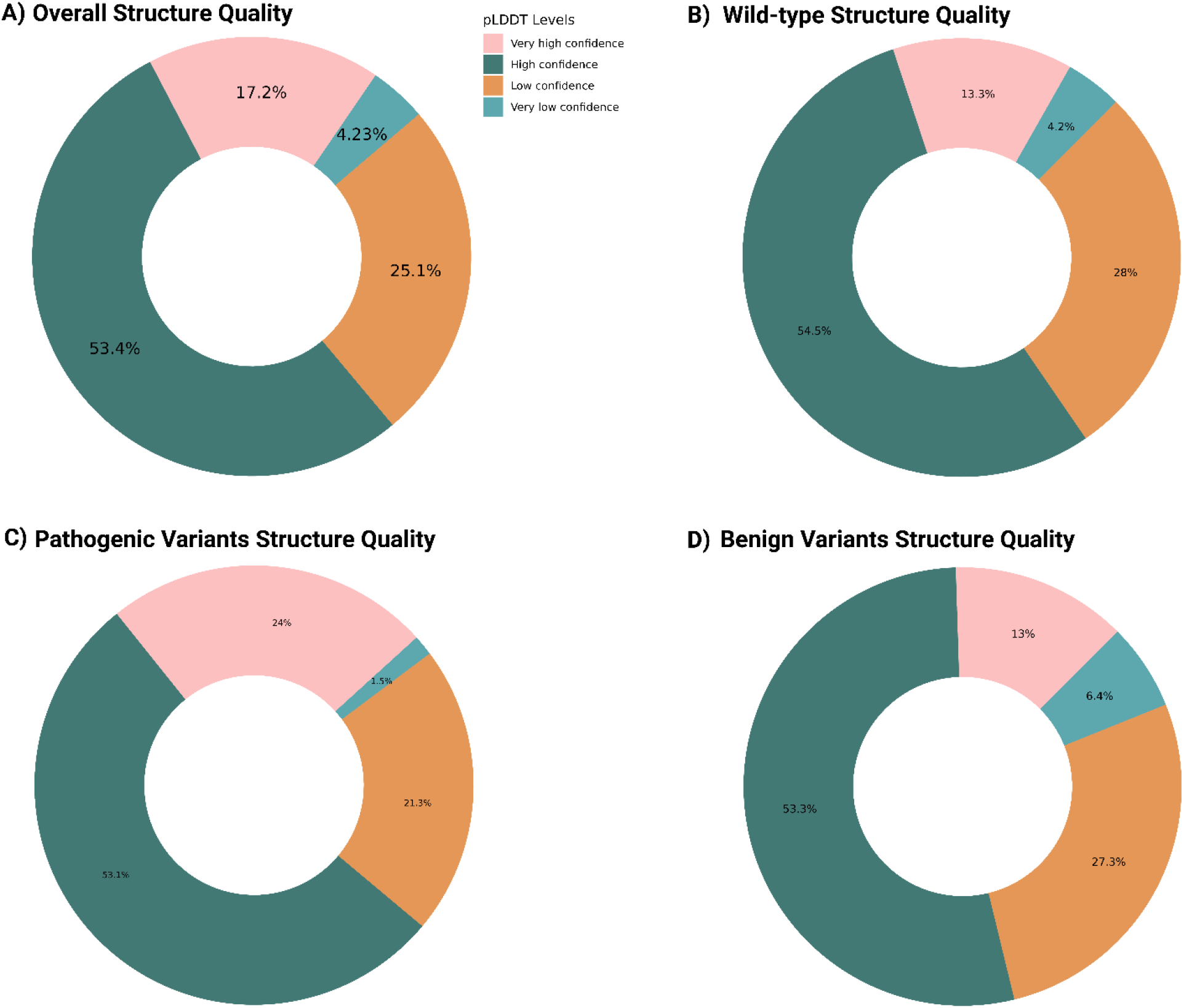
Distribution of pLDDT scores across the predicted structures. **A)** Overall quality of the predicted wild-type and variant (pathogenic and benign) structures across four pLDDT levels. **B)** Quality of the predicted wild-type protein structures. **C)** Quality of the predicted pathogenic variant structures. **D)** Quality of the predicted benign variant structures.

Additionally, we assessed the prediction quality of the predicted wild-type, pathogenic, and benign structures individually (Figures 2B/2C/2D). We found that models belonging to pathogenic missense variants had a high proportion of structures predicted with very high confidence (24%) compared to wild-type (13.3%) and benign variants (13%). A similar trend was observed in the very low-confidence category, where only 1.5% of the models belonging to pathogenic missense variants had pLDDT values <50, in contrast to 4.2% of the wild-type and 6.4% of the benign structures (Figures 2B/2C/2D). Meanwhile, the proportion of structures predicted with high confidence (70 ≤ pLDDT < 90) was relatively consistent across all three categories, exceeding 50%.

### Data access and retrieval from the DeltaMut Webserver

AlphaFold2 generates five structural models per protein, each provided in both relaxed and unrelaxed forms. The relaxed models undergo an additional energy minimization step to resolve atomic clashes, whereas the unrelaxed models represent the raw neural network prediction before the relaxation step. Both the relaxed and unrelaxed models are freely downloadable in PDB format from the DeltaMut web server without restrictions. In DeltaMut, wild-type structures are named using the corresponding UniProt accession ID of the modeled protein. For the variant structures, the file name consists of the UniProt accession ID concatenated with amino acid substitution in the following format *<reference amino acid, position, alternative amino acid>* (e.g., P38398_N40Y).

Additional files generated during the prediction process, such as *features*.*pkl*, which contains the output of the MSA construction phase, and *results_model*.*pkl* of the best-ranked model are currently not available for download via the web server due to storage limitations. However, users can freely obtain these files through the HUN-REN ARP repository, as described in https://github.com/erdaqorri/DeltaMut-Database.

### Querying the DeltaMut Webserver

The DeltaMut database can be queried using various inputs such as UniProt accession IDs, protein or gene names, or pathway-related keywords. For instance, searching for “Mismatch repair” will retrieve all genes linked to this function included in the database (Figure 3A).

**Figure 3:**
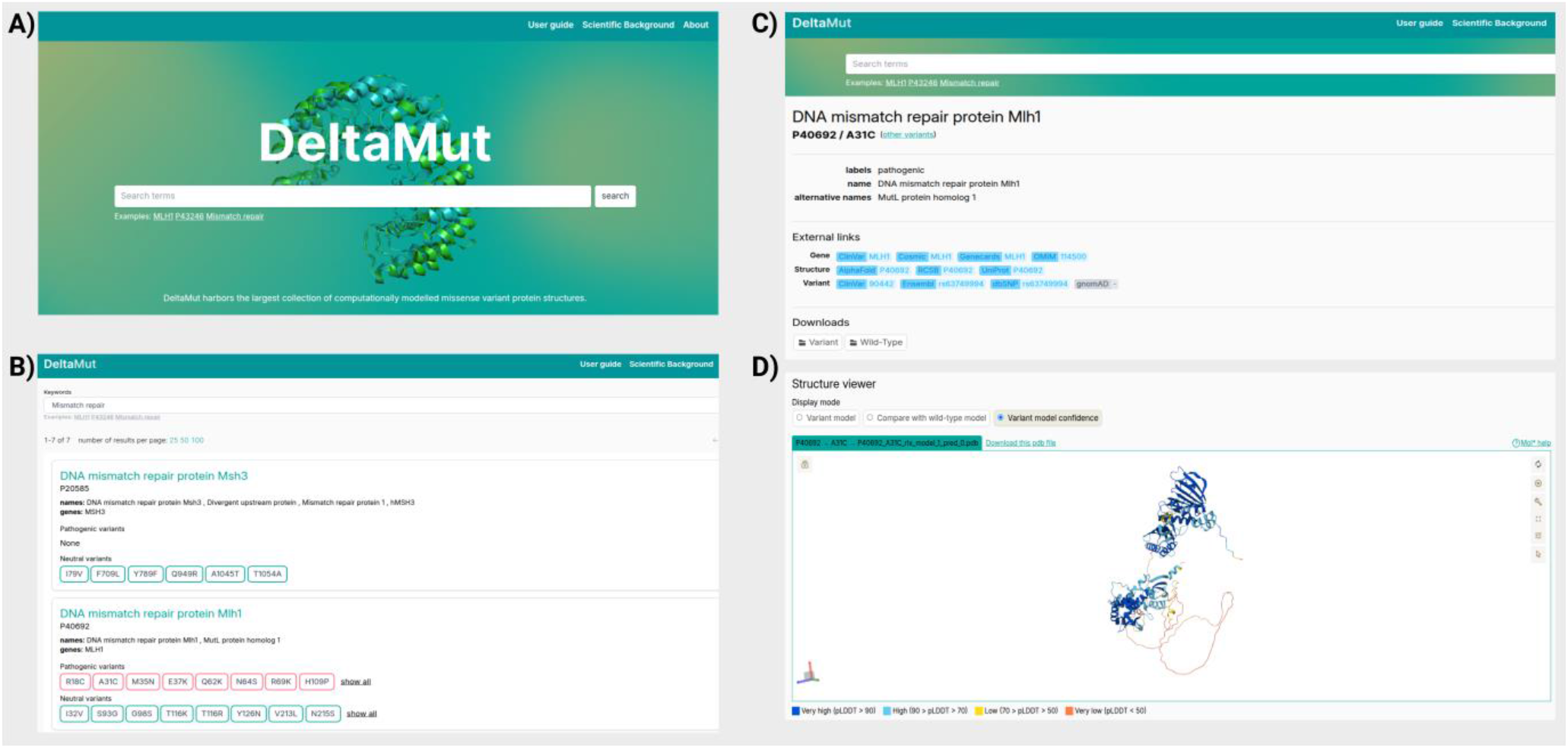
Screen captures demonstrating the DeltaMut webserver and its core functionalities. **A)** The homepage, which allows users to query the database using various keywords. **B)** Query search results page. **C)** Gene-specific page providing detailed information on the selected gene, links to external databases, and options for downloading structural data. **D)** Interactive structure viewer page for visualizing 3D protein models.

After submitting a query, users are directed to a dedicated webpage for the selected protein, displaying essential information such as the UniProt accession ID, gene name, and the corresponding list of pathogenic and neutral variants, each presented in individual boxes, color-coded in red and green, respectively (Figure 3B). Selecting a specific variant redirects to a new webpage, in which three main sections can be identified: *External Links, Downloads, and Structure Viewer*. The *External Links* section provides direct access to variant-specific entries (when available) across three different database categories: *Gene*: ClinGen (Rehm et al. 2015), ClinVar (Landrum et al. 2016), Cosmic (Sondka et al. 2024), GeneCards (Stelzer et al. 2016), and OMIM (Amberger et al. 2015); *Structure*: AlphaFold Protein Structure Database (Varadi et al. 2022), RSCB PDB Database (Berman, Henrick, and Nakamura 2003), and UniProt (Consortium 2024); and *Variant*: ClinVar (Landrum et al. 2016), Ensembl (Dyer et al. 2024), dbSNP (Sherry et al. 2001), and gnomAD (Karczewski et al. 2020).

The *Downloads* section provides easy access to download both the relaxed and unrelaxed AlphaFold2 models for the wild-type and variant structures in PDB format. If the “Download All” option is selected, the models are provided in a zip file format (Figure 3C).

Lastly, the *Structure Viewer* section allows users to visually explore the predicted structure of the variant, compare it in real-time with the wild-type model through superimposition, and assess the prediction confidence based on the assigned pLDDT scores (Figure 3D). To further enhance user experience, the structure is displayed by default with a zoomed-in view of the mutation site highlighted in red, and a quick access link to the Mol*Viewer documentation is provided for additional assistance.

## Conclusion and Future Perspectives

Here we present DeltaMut, the first publicly available database of large-scale AlphaFold2-predicted structures for missense variants. DeltaMut includes over 77,000 protein structures, including 65,612 missense variant models, with 70.5% of the structures predicted at high confidence. By providing both variants and corresponding wild-type structures, DeltaMut fills in a critical gap in structural bioinformatics. While its primary application lies in exploring the potential of variant structures in improving missense variant effect prediction approaches, it can also support *in silico* drug design and can potentially be integrated into omics analysis to provide structural insights. With unrestricted bulk downloads, interactive visualization tools, and disease-relevant protein coverage, DeltaMut offers a unique resource for both computational and experimental researchers. Future updates to DeltaMut will expand coverage to ClinVar variants, including proteins longer than 4000 amino acids and will offer direct webserver access to additional AlphaFold2 output files.

## Availability of Supporting Data

The predicted PDB files are available for download directly from the DeltaMut website, while all additional files generated during the prediction process can be accessed from the ARP repository. Detailed information on how the data can be retrieved from the ARP repository can be found in the following GitHub repository: https://github.com/erdaqorri/DeltaMut-Database/tree/main#deltamut-protein-structure-database-acess-and-download

## Declaration of Generative AI and AI-assisted Technologies in the Writing Process

ChatGPT was utilized by the authors during the preparation of this work to enhance the language quality of the manuscript. The authors state that the AI-edited content was reviewed and edited as needed and therefore take full responsibility for the content included for publication.

## CRediT Authorship Contribution Statement

**Erda Qorri**: Conceptualization, Methodology, Software, Data curation, Validation, Formal analysis, Writing – original draft, Writing – review & editing. **Krisztián Ádám**: Data curation, Software, Writing – review & editing. **Bertalan Takács**: Software, Data curation, Visualization, Writing – review & editing. **Sára Shemesh**: Visualization and Writing. **Dénes Buzafalvi**: Formal analysis, Methodology, Validation, Writing – review & editing. **Valentin Varga**: Visualization, Data curation, Writing – review & editing. **Emese Pekker**: Visualization, Writing – review & editing. **Lajos Pintér**: Conceptualization, Methodology, Project administration, Resources, Supervision. **Zoltán Hegedűs**: Validation, Writing – review & editing. **Lajos Haracska**: Conceptualization, Funding acquisition, Resources, Supervision, Writing – review & editing.

## Acknowledgements

We would like to acknowledge the provider of the supercomputing service, Digital Government Development and Project Management Ltd. (DKF) for allowing continued access to Komondor HPC. The authors would like to thank the system administrators assigned to this project, particularly Attila Debreczeni, Zoltán Kiss, and Dr. Tamás István as well as Erzsébet Horváth for overseeing the project. Additionally, we thank Péter Pállinger from the SZTAKI Department of Distributed Systems (DSD) for providing technical assistance during the data transfer to the ARP repository. Moreover, the authors would like to thank Gabriella Tick for proofreading the manuscript.

This work was supported by the National Research, Development, and Innovation Office (2023-1-1-1-PIACI-FÓKUSZ-2024-00029, 2024-1.1.1-KKV_FÓKUSZ-2024-00019, and 2025-1.1.2-GYORSíTÓSÁV-2025-00038).

## References

Adzhubei, Ivan, Daniel M. Jordan, and Shamil R. Sunyaev. 2013. Predicting Functional Effect of Human Missense Mutations Using PolyPhen-2. Current Protocols in Human Genetics. 10.1002/0471142905.hg0720s76.

Amberger, Joanna S., Carol A. Bocchini, François Schiettecatte, Alan F. Scott, and Ada Hamosh. 2015. “OMIM.Org: Online Mendelian Inheritance in Man (OMIM®), an Online Catalog of Human Genes and Genetic Disorders.” Nucleic Acids Research 43 (D1). 10.1093/nar/gku1205.

Baek, Minkyung, Frank DiMaio, Ivan Anishchenko, Justas Dauparas, Sergey Ovchinnikov, Gyu Rie Lee, Jue Wang, et al. 2021. “Accurate Prediction of Protein Structures and Interactions Using a Three-Track Neural Network.” Science 373 (6557). https://doi.org/10.1126/science.abj8754.

Berman, Helen, Kim Henrick, and Haruki Nakamura. 2003. “Announcing the Worldwide Protein Data Bank.” Nature Structural Biology. 10.1038/nsb1203-980.

Bromberg, Yana, and Burkhard Rost. 2007. “SNAP: Predict Effect of Non-Synonymous Polymorphisms on Function.” Nucleic Acids Research 35 (11). 10.1093/nar/gkm238.

Cheng, Jun, Guido Novati, Joshua Pan, Clare Bycroft, Akvilė Žemgulytė, Taylor Applebaum, Alexander Pritzel, et al. 2023. “Accurate Proteome-Wide Missense Variant Effect Prediction with AlphaMissense.” Science. 10.1126/science.adg7492.

Chris McCord. n.d. “Phoenix Framework.” https://phoenixframework.org/.

Consortium, The UniProt. 2024. “UniProt: The Universal Protein Knowledgebase in 2025.” Nucleic Acids Research 53 (D1): D609–17. 10.1093/nar/gkae1010.

Dyer, Sarah C, Olanrewaju Austine-Orimoloye, Andrey G Azov, Matthieu Barba, If Barnes, Vianey Paola Barrera-Enriquez, Arne Becker, et al. 2024. “Ensembl 2025.” Nucleic Acids Research 53 (D1): D948–57. 10.1093/nar/gkae1071.

Elixir Core Team. n.d. “Elixir.” https://elixir-lang.org/.

Federici, Giulia, and Silvia Soddu. 2020. “Variants of Uncertain Significance in the Era of High-Throughput Genome Sequencing: A Lesson from Breast and Ovary Cancers.” Journal of Experimental and Clinical Cancer Research 39 (1): 1–12. 10.1186/s13046-020-01554-6.

Frazer, Jonathan, Pascal Notin, Mafalda Dias, Aidan Gomez, Joseph K. Min, Kelly Brock, Yarin Gal, and Debora S. Marks. 2021. “Disease Variant Prediction with Deep Generative Models of Evolutionary Data.” Nature 599 (7883). https://doi.org/10.1038/s41586-021-04043-8.

Hatano, Yuya, Tomohiko Ishihara, and Osamu Onodera. 2023. “Accuracy of a Machine Learning Method Based on Structural and Locational Information from AlphaFold2 for Predicting the Pathogenicity of TARDBP and FUS Gene Variants in ALS.” BMC Bioinformatics 24 (1). 10.1186/s12859-023-05338-5.

Hipp, Richard D. 2020. “SQLite.” https://www.sqlite.org/index.html.

Ioannidis, Nilah M., Joseph H. Rothstein, Vikas Pejaver, Sumit Middha, Shannon K. McDonnell, Saurabh Baheti, Anthony Musolf, et al. 2016. “REVEL: An Ensemble Method for Predicting the Pathogenicity of Rare Missense Variants.” American Journal of Human Genetics 99 (4): 877–85. 10.1016/j.ajhg.2016.08.016.

Jumper, John, Richard Evans, Alexander Pritzel, Tim Green, Michael Figurnov, Olaf Ronneberger, Kathryn Tunyasuvunakool, et al. 2021. “Highly Accurate Protein Structure Prediction with AlphaFold.” Nature 596 (7873). 10.1038/s41586-021-03819-2.

Karczewski, Konrad J, Laurent C Francioli, Grace Tiao, Beryl B Cummings, Jessica Alföldi, Qingbo Wang, Ryan L Collins, et al. 2020. “The Mutational Constraint Spectrum Quantified from Variation in 141,456 Humans.” Nature 581 (7809): 434–43. 10.1038/s41586-020-2308-7.

Landrum, Melissa J., Jennifer M. Lee, Mark Benson, Garth Brown, Chen Chao, Shanmuga Chitipiralla, Baoshan Gu, et al. 2016. “ClinVar: Public Archive of Interpretations of Clinically Relevant Variants.” Nucleic Acids Research 44 (D1): D862–68. 10.1093/nar/gkv1222.

Landrum, Melissa J., Jennifer M. Lee, George R. Riley, Wonhee Jang, Wendy S. Rubinstein, Deanna M. Church, and Donna R. Maglott. 2014. “ClinVar: Public Archive of Relationships among Sequence Variation and Human Phenotype.” Nucleic Acids Research 42 (D1). 10.1093/nar/gkt1113.

Ng, Pauline C., and Steven Henikoff. 2002. “Accounting for Human Polymorphisms Predicted to Affect Protein Function.” Genome Research 12 (3): 436–46. 10.1101/gr.212802.

Ozkan, Selen, Natàlia Padilla, and Xavier de la Cruz. 2024. “QAFI: A Novel Method for Quantitative Estimation of Missense Variant Impact Using Protein-Specific Predictors and Ensemble Learning.” Human Genetics. 10.1007/s00439-024-02692-z.

Rehm, Heidi L., Jonathan S. Berg, Lisa D. Brooks, Carlos D. Bustamante, James P. Evans, Melissa J. Landrum, David H. Ledbetter, et al. 2015. “ClinGen — The Clinical Genome Resource.” New England Journal of Medicine 372 (23). 10.1056/nejmsr1406261.

Reza, Mahjerin Nasrin, Nadim Ferdous, Md Tabassum Hossain Emon, Md Shariful Islam, A. K.M. Mohiuddin, and Mohammad Uzzal Hossain. 2021. “Pathogenic Genetic Variants from Highly Connected Cancer Susceptibility Genes Confer the Loss of Structural Stability.” Scientific Reports 11 (1). 10.1038/s41598-021-98547-y.

Schmidt, Axel, Sebastian Röner, Karola Mai, Hannah Klinkhammer, Martin Kircher, and Kerstin U. Ludwig. 2023. “Predicting the Pathogenicity of Missense Variants Using Features Derived from AlphaFold2.” Bioinformatics 39 (5). 10.1093/bioinformatics/btad280.

Sehnal, David, Sebastian Bittrich, Mandar Deshpande, Radka Svobodová, Karel Berka, Václav Bazgier, Sameer Velankar, Stephen K. Burley, Jaroslav Koča, and Alexander S. Rose. 2021. “Mol∗Viewer: Modern Web App for 3D Visualization and Analysis of Large Biomolecular Structures.” Nucleic Acids Research 49 (W1). 10.1093/nar/gkab314.

Sherry, S. T., M. H. Ward, M. Kholodov, J. Baker, L. Phan, E. M. Smigielski, and K. Sirotkin. 2001. “DbSNP: The NCBI Database of Genetic Variation.” Nucleic Acids Research 29 (1). 10.1093/nar/29.1.308.

Sondka, Zbyslaw, Nidhi Bindal Dhir, Denise Carvalho-Silva, Steven Jupe Madhumita, Karen McLaren, Mike Starkey, et al. 2024. “COSMIC: A Curated Database of Somatic Variants and Clinical Data for Cancer.” Nucleic Acids Research 52 (D1). 10.1093/nar/gkad986.

Stelzer, Gil, Naomi Rosen, Inbar Plaschkes, Shahar Zimmerman, Michal Twik, Simon Fishilevich, Tsippi Iny Stein, et al. 2016. “The GeneCards Suite: From Gene Data Mining to Disease Genome Sequence Analyses.” Current Protocols in Bioinformatics 2016. 10.1002/cpbi.5.

“UniProt: The Universal Protein Knowledgebase in 2025.” 2025. Nucleic Acids Research 53 (D1): D609–17. 10.1093/nar/gkae1010.

Varadi, Mihaly, Stephen Anyango, Mandar Deshpande, Sreenath Nair, Cindy Natassia, Galabina Yordanova, David Yuan, et al. 2022. “AlphaFold Protein Structure Database: Massively Expanding the Structural Coverage of Protein-Sequence Space with High-Accuracy Models.” Nucleic Acids Research 50 (D1). 10.1093/nar/gkab1061.

Zhong, Bozitao, Xiaoming Su, Minhua Wen, Sicheng Zuo, Liang Hong, and James Lin. 2022. “ParaFold: Paralleling AlphaFold for Large-Scale Predictions.” In ACM International Conference Proceeding Series. 10.1145/3503470.3503471.

